# Turnover of MCMV-expanded CD8^+^ T-cells is similar to that of memory phenotype T-cells and independent of the magnitude of the response

**DOI:** 10.1101/2021.11.03.467089

**Authors:** Mariona Baliu-Piqué, Julia Drylewicz, Xiaoyan Zheng, Lisa Borkner, Arpit C. Swain, Sigrid Otto, Rob J. de Boer, Kiki Tesselaar, Luka Cicin-Sain, José A.M. Borghans

**Affiliations:** Center for Translational Immunology, University Medical Center Utrecht, Utrecht, The Netherlands; Immunotherapy Manufacturing Center, Galaria-Sergas, Santiago de Compostela, Spain; Department of Viral Immunology, Helmholtz Centre for Infection Research, Braunschweig, Germany; Theoretical Biology, Utrecht University, Utrecht, The Netherlands; German Centre for Infection Research (DZIF), Hannover-Braunschweig partner site

## Abstract

The potential of memory T-cells to provide protection against re-infection is beyond question. Yet, it remains debated whether long-term T-cell memory is due to long-lived memory cells. There is ample evidence that blood-derived memory phenotype CD8^+^ T-cells maintain themselves through cell division, rather than through longevity of individual cells. It has recently been proposed, however, that there may be heterogeneity in the lifespans of memory T-cells, depending on factors such as exposure to cognate antigen. Cytomegalovirus (CMV) infection induces not only conventional, contracting T-cell responses, but also inflationary CD8^+^ T-cell responses, which are maintained at unusually high numbers, and are even thought to continue to expand over time. It has been proposed that such inflating T-cell responses result from the accumulation of relatively long-lived CMV-specific memory CD8^+^ T-cells. Using *in vivo* deuterium labelling and mathematical modelling, we found that the average production rates and expected lifespans of mouse CMV-specific CD8^+^ T-cells are very similar to those of bulk memory-phenotype CD8^+^ T-cells. Even CMV-specific inflationary CD8^+^ T-cell responses that differ three-fold in size, were found to turn over at similar rates.

## Introduction

Memory CD8^+^ T-cells are a crucial component of the adaptive immune response to viruses. Antigen-specific memory CD8^+^ T-cells convey immune protection against viral infections that may last for long periods of time, sometimes even life-long. There is ample evidence that memory T-cells isolated from the blood and lymph nodes are relatively short-lived. Their lifespan is much shorter than that of naive T-cells, and far shorter than the long-term immune protection they convey (1–10). Memory T-cell populations are heterogeneous, both phenotypically and functionally. They consist of phenotypically defined subpopulations, such as central memory (T_CM_) and effector memory (T_EM_) T-cells, and of subsets that differ in terms of exposure to their cognate antigen. *In vivo* deuterium labelling studies have shown that different subsets of memory T-cells can have different kinetics. CD4^+^ T_EM_ cells were shown to have shorter lifespans than T_CM_ cells (10), and yellow fever virus (YFV)-specific memory T-cells generated by vaccination, which can persist for years, were found to have longer lifespans than bulk memory-phenotype cells (11).

Cytomegalovirus (CMV) infection is a persistent, chronic infection, which, in contrast to YFV-vaccination, results in continual antigen presentation. CMV is under constant immune surveillance, and triggers ongoing CD8^+^ T-cell responses that provide effective viral control for long periods of time. A hallmark of the CD8^+^ T-cell response to CMV infection is the steady maintenance or accumulation of large populations of virus-specific effector CD8^+^ T-cells over time, a phenomenon termed memory inflation (12). Expanded CD8^+^ T-cell populations specific for unique CMV epitopes can become extraordinarily large, composing up to 20% of the total memory CD8^+^ T-cell pool (13–18). The fact that inflationary CD8^+^ T-cell responses are maintained at such high levels, has led to the hypothesis that T-cell inflation may arise from the accumulation of relatively long-lived CMV-specific memory CD8^+^ T-cells (19).

Here we addressed this hypothesis using *in vivo* deuterium labelling and mathematical modelling, the state-of-the-art techniques to quantify lymphocyte turnover, in the setting of murine cytomegalovirus (MCMV) infection, a relevant experimental model to study memory T-cell inflation (14). In contrast to the postulated hypothesis that CMV-specific T-cells may have extended lifespans, we found no significant difference in the expected lifespans of MCMV-specific CD8^+^ T-cells and bulk memory-phenotype CD8^+^ T-cells. Using recombinant viruses inducing inflationary CD8^+^ T-cell responses of different magnitudes, we found that MCMV-specific T-cells composing small and large inflationary T-cell responses had very similar turnover rates.

## Material and Methods

### Mice

129S2/SvPas Crl (129/Sv) mice were purchased from Charles River (Sulzfeld, Germany). Mice were housed and handled in accordance with good animal practice as defined by the Federation of Laboratory Animal Science Associations and the national animal welfare body “Die Gesellschaft für Versuchstierkunde”/Society of Laboratory Animals. All animal experiments were approved by the responsible state office (Lower Saxony State Office of Consumer Protection and Food Safety, Germany; permit NO. 33.19-42502-04-15/1836 and by the Animal Experiments Committee of Utrecht University, IVD Utrecht, The Netherlands; DEC AVD115002016714).

### Viruses

Bacterial artificial chromosome (BAC)-derived recombinant viruses MCMV^ie2SL^ and MCMV^M45SL^ were generated and propagated as described previously (18), and the recombinant virus MCMV^ie2KNL^ was generated and propagated as described in Borkner *et al*. (20).

### In vivo infection

Female 8-wk-old mice were infected with 2×10^5^ PFU MCMV^ie2SL^ (N=40), MCMV^M45SL^ (N=37) or MCMV^ie2KNL^ (N=41) and housed in specific pathogen-free conditions throughout the experiment. Non-infected, sex- and age-matched mice were used as controls (N=10).

### Stable isotope labelling

120 days after MCMV infection, mice received 8% deuterated water (99.8% ^2^H_2_O, Cambridge Isotope Laboratories) in their drinking water for 28 days. At day 4, mice were given an intra peritoneal (i.p.) boost injection of 15ml/kg ^2^H_2_O in phosphate-buffered saline (PBS). To determine deuterium enrichment in the body water, EDTA-plasma was collected during the up- and down-labelling phase, and was frozen and stored at −80°C until analysis.

### Sampling and cell preparation

Spleen, thymus and blood were isolated at different time points during and after label administration. Blood was collected in EDTA tubes. Single cell suspensions from blood, spleen and thymus were obtained as described previously (21).

### Flow cytometry and cell sorting

To determine the fraction of SSIEFARL (SL) and KCSRNRQYL (KNL) epitope-specific T-cells, single cell suspensions from blood and spleen were stained with allophycocyanin-conjugated SL-K^b^ or KNL-D^b^ tetramers (Tet) for 15 min at 4°C. Samples were further stained for 30 min at 4°C with anti-CD3-APC-eFluor780 (clone 17A2; eBioscience), anti-CD3e-FITC (clone 145-2C11; BD Phamingen), anti-CD3-V500 (clone 500A2; BD), anti-CD4-Pacific Blue (clone GK1.5; BioLegend), anti-CD4-Brilliant violet 650 (clone GK1.5; BD Harizon), anti-CD4-APC-H7 (GK1.5; BD), anti-CD8a-PerCP/Cy5.5 (clone 53-6.7; BioLegend), anti-CD8a-BV786 (clone 53-6.7; BD), anti-CD44-Alexa Fluor700 (clone IM7; BioLegend), anti-CD44-Alexa Fluor450 (IM7; eBioscience), anti-CD62L-eVolve 605 (clone MEL-14; eBioscience), anti-CD62L-FITC (MEL-14; eBioscience), anti-CD127-PE/Cy7 (clone A7R34; Biolegend) monoclonal antibodies. For intracellular staining, cells were subsequently fixed for 20 min at room temperature with 100μl fixation/permeabilisation buffer of the FoxP3 Transcription factor staining set (eBioscience), permeabilised for 15 min at room temperature in 100μl permeabilisation buffer (eBioscience), and stained with Ki67-PE (clone 16A8, Biolegend) in 100μl permeabilisation buffer for 30 min at room temperature. Cells were analysed on an LSR-Fortessa flow cytometer using FACS Diva software (BD Biosciences) and FlowJo software (version 9.8.3). For infected mice, tetramer-positive (CD3^+^CD8^+^Tet^+^), and tetramer-negative naive (T_N_, CD3^+^CD8^+^Tet^−^CD62L^+^CD44^−^), central memory (T_CM_, CD3^+^CD8^+^Tet^−^CD62L^+^CD44^+^), and effector memory (T_EM_, CD3^+^CD8^+^Tet^−^CD62L^−^CD44^+^) T-cells were sorted from spleen on a FACS Aria-II SORP (BD Biosciences), FACS Aria III (BD Biosciences) or MoFlo XDP cell sorter (Sup. Figure 1). For uninfected mice, T_N_, T_CM_, and T_EM_ cells were sorted from spleen on a FACS Aria III (BD Biosciences).

### DNA isolation

Genomic DNA was isolated from total thymocytes and from sorted T-cell subsets from MCMV-infected and uninfected mice according to the manufacturer’s instructions using the NucleoSpin Blood QuickPure kit (MACHEREY-NAGEL), and stored at −20°C until further analysis.

### Measurement of ^2^H_2_O enrichment in body water and DNA

Deuterium enrichment in plasma and DNA was analysed by gas-chromatography/mass-spectrometry (GC/MS) using an Agilent 5973/6890 GC/MS system (Agilent Technologies). Plasma was derivatized to acetylene (C_2_H_2_) as previously described (5). The derivative was injected into the GC/MS equipped with a PoraPLOT Q 25×0.32 column (Varian), and measured in SIM mode monitoring ions m/z 26 (M+0) and m/z 27 (M+1). From the ratio of ions, plasma deuterium enrichment was calculated by calibration against standard samples of known enrichment. DNA obtained from sorted lymphocytes and granulocytes was hydrolysed to deoxy-ribonucleotides and derivatized to penta-fluoro-triacetate (PFTA) (5). The derivative was injected into the GC/MS equipped with a DB-17 column (Agilent Technologies) and measured in SIM mode monitoring ions m/z 435 (M+0), and m/z 436 (M+1). From the ratio of ions, we calculated DNA deuterium enrichment by calibration against deoxyadenosine standards of known enrichment, as previously described (22).

### Mathematical modelling of T-cell dynamics

We deduced the dynamics of tetramer-negative naive (T_N_), tetramer-negative central memory (T_CM_), tetramer-negative effector memory (T_EM_), and tetramer-negative total memory (T_M_, calculated as the weighted average of T_CM_ and T_EM_ cells) CD8^+^ T-cells, and of tetramer-positive (Tet^+^) CD8^+^ T-cells from the deuterium labelling data using previously published mathematical models (3, 23). In brief, to monitor the changing levels of deuterium (^2^H) in body water over the course of the experiment, a simple label enrichment/decay curve was fitted to the ^2^H enrichment in plasma (3):

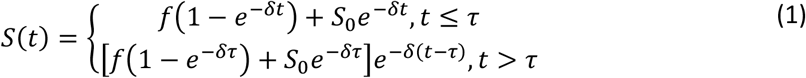

where *S(t)* is the fraction of deuterium in plasma at time *t* (in days), *f* is the predicted plateau value of deuterium enrichment in the plasma, *δ* is the turnover rate of body water per day, *S_0_* is the plasma enrichment level attained due to the i.p. ^2^H_2_O boost, and ^2^H_2_O administration was stopped at *τ* = 28 days. The level of label incorporation in the different cell subsets was described by:

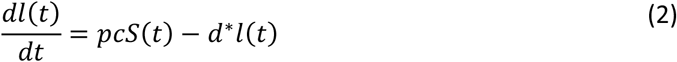

Here, *l(t)* is the fraction of labelled DNA in the cell subset, *p* is the average (per capita) production rate of the cells, *d*\* is the average (per capita) rate at which *labelled* cells are lost (which need not be equal to the average loss rate of cells in the population (23)) and *c* is an amplification factor, which accounts for the multiple hydrogen atoms that can be replaced by deuterium (see (3)). To estimate the value of *c*, we first fitted Equation (2) for a kinetically homogeneous population (*p* = *d*\*) to the level of deuterium enrichment in the DNA of total thymocytes, as they are known to have a high turnover rate (5). The resulting estimated value of *c* was subsequently fixed to estimate the turnover rates of T_N_, T_CM_, T_EM_, and T_M_ and Tet^+^ CD8^+^ T-cells. The best fits to the plasma and thymocyte data are shown in Sup. Figure 2 (see Sup. Table 1 for the estimated parameter values). When modelling the deuterium enrichment levels of T_N_ cells, a time delay Δ was introduced between T-cell production in the thymus and the appearance of labelled DNA in naive T-cells in the spleen, based on previous observations (5). This was done by incorporating a delayed labelling curve of the deuterium enrichment in plasma (i.e., *S*(*t*-Δ)) in Equation (2) when fitting the dynamics of T_N_ cells.

To estimate the rate of change in cell numbers, *r*, in the T_N_, T_CM_, T_EM_, T_M_, and Tet^+^ T-cell populations, we used a simple exponential growth/decay model, 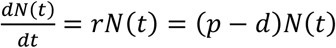, which we fitted to the cell number data from the start of the experiment (i.e. 120 days p.i.) until 550 days later (see Table 2 and Sup. Figure 5). Results were very similar when *r* was estimated based on cell numbers during the first 140 days of the experiment. Based on the resulting value of *r* and the estimated value of *p* from Equation (2), the cellular loss rates, *p* – *r*, were calculated (see Table 2). The expected lifespans of cells can be calculated as the inverse of their average loss rates, i.e., as 1/*d*.

**Table 1.** Average production rates (*p*) and loss rates of labelled cells (*d\**) in MCMV-infected mice. Estimated parameters and their corresponding 95% confidence intervals are shown. For T_N_ cells, we report the best fits of the model with *p* = *d*\* as allowing for different values of *p* and *d*\* did not significantly improve the fit to the data (P-values of F-test=1 for MCMV^ie2SL^, P-values of F-test=0.09 for MCMV^ie2KNL^, and P-values of F-test=0.54 for MCMV^m45SL^).

| Average production rate per day ($p$) | | | |
| --- | --- | --- | --- |
| CD8<sup>+</sup> T-cell subset | MCMV<sup>ie2SL</sup> | MCMV<sup>ie2KNL</sup> | MCMV<sup>m45SL</sup> |
| $T_N$ | 0.0090 (0.0084;0.0095) | 0.0095 (0.0085;0.0107) | 0.0077 (0.0068;0.0089) |
| $T_{CM}$ | 0.0132 (0.0114;0.0154) | 0.0151 (0.0133;0.0177) | 0.0139 (0.0126;0.0153) |
| $T_{EM}$ | 0.0204 (0.0184;0.0229) | 0.0229 (0.0203;0.0270) | 0.0205 (0.0192;0.0221) |
| $T_M$ | 0.0169 (0.0149;0.0188) | 0.0192 (0.0172;0.0219) | 0.0182 (0.0171;0.0195) |
| $Tet^+$ | 0.0169 (0.0158;0.0182) | 0.0195 (0.0175;0.0222) | 0.0161 (0.0147;0.0176) |
| Average loss rate of labelled cells per day ($d^*$) | | | |
| CD8<sup>+</sup> T-cell subset | MCMV<sup>ie2SL</sup> | MCMV<sup>ie2KNL</sup> | MCMV<sup>m45SL</sup> |
| $T_N$ | 0.0090 (0.0084;0.0095) | 0.0095 (0.0085;0.0107) | 0.0077 (0.0068;0.0089) |
| $T_{CM}$ | 0.0131 (0.0111;0.0160) | 0.0113 (0.0093;0.0142) | 0.0115 (0.0102;0.0130) |
| $T_{EM}$ | 0.0146 (0.0130;0.0163) | 0.0153 (0.0134;0.0180) | 0.0157 (0.0143;0.0170) |
| $T_M$ | 0.0141 (0.0124;0.0158) | 0.0135 (0.0118;0.0160) | 0.0146 (0.0135;0.0161) |
| $Tet^+$ | 0.0116 (0.0104;0.0128) | 0.0120 (0.0100;0.0143) | 0.0126 (0.0113;0.0140) |

**Table 2.** Average population growth rates (*r*) and average loss rates (*d*) in MCMV-infected mice. Loss rates *d* were calculated using the estimated average production rates *p* (from the deuterium labelling experiments, Table 1) and the estimated overall growth/decay rates *r* of the specific T-cell populations (followed over 550 days, Sup. Figure 5), see Methods. Overall growth/decay rates *r* of the specific T-cell populations were estimated by simultaneously estimating each population size at the start of the experiment (i.e. 120 days p.i.), N(0), of which the values are given in Sup. Table 2. Estimated growth rates (r) are reported with their corresponding 95% confidence intervals in brackets. Because the average loss rates *d* were calculated based on other parameters, they are given without 95% confidence intervals.

| Average growth/decay rate of the cell population per day ($r$) | | | |
| --- | --- | --- | --- |
| CD8<sup>+</sup> T-cell subset | MCMV<sup>ie2SL</sup> | MCMV<sup>ie2KNL</sup> | MCMV<sup>m45SL</sup> |
| T<sub>N</sub> | -0.00093 (-0.0028;0.00015) | -0.00094 (-0.0039;0.00065) | -0.0016 (-0.004;-0.0004) |
| T<sub>CM</sub> | 0.0014 (0.00032;0.0024) | 0.0017 (-0.0028;0.0027) | 0.0027 (0.0007;0.0044) |
| T<sub>EM</sub> | 0.0026 (0.0013;0.0037) | 0.0040 (0.00025;0.0068) | 0.0015 (0.00014;0.0028) |
| T<sub>M</sub> | 0.0020 (0.000895;0.0029) | 0.0030 (-0.00033;0.0043) | 0.0021 (0.00083;0.0033) |
| Tet<sup>+</sup> | 0.0032 (0.0021;0.0045) | 0.001 (-0.0021;0.008) | 0.0014 (-0.00025;0.0029) |
| Calculated average loss rate per day ($d$) | | | |
| CD8<sup>+</sup> T-cell subset | MCMV<sup>ie2SL</sup> | MCMV<sup>ie2KNL</sup> | MCMV<sup>m45SL</sup> |
| T<sub>N</sub> | 0.0099 | 0.0105 | 0.00939 |
| T<sub>CM</sub> | 0.0118 | 0.0134 | 0.0112 |
| T<sub>EM</sub> | 0.0178 | 0.0189 | 0.0190 |
| T<sub>M</sub> | 0.0149 | 0.0162 | 0.0161 |
| Tet<sup>+</sup> | 0.0137 | 0.0185 | 0.0147 |

Best fits to the labelling and cell number data were determined by minimizing the sum of squared residuals (SSR) using the R function modCost() of the FME package (24). The fractions of labelled DNA, x, were transformed using the function arcsin(sqrt(x)) before the fitting procedure. Fitting the cell number data yielded estimates for the initial cell number at the start of the experiment at 120 days p.i, *N*(0), and the exponential growth rate, *r*. The 95% confidence intervals (CI) on the estimated parameters for both labelling and cell number data were determined using a bootstrap method where the data points were resampled 500 times. Fitting the exponential growth/decay model to these 500 data samples, yielded 500 bootstrap trajectories. The 95% CI trajectories for the cell numbers were calculated by taking the 95% CI of these 500 bootstraps at each time point.

### Statistical analysis

Statistical analyses were performed using GraphPad Prism. Comparisons between two and more groups were performed using Kruskal-Wallis and Dunn’s multiple comparisons test. P-values <0.05 were considered significant.

## Results

### Induction of inflationary CD8^+^ T-cell responses of different magnitude and phenotype

To study the kinetics of MCMV-specific and inflationary CD8^+^ T-cell responses during the memory phase of MCMV infection, we made use of 129/Sv mice, to benefit from the well-defined H^2b^ MHC-I haplotype and the well-characterized arrays of epitopes associated with it, while circumventing a protective dominant role of NK cells in controlling the infection (18, 25). The avidity of the viral epitope together with its context of gene expression define the kinetics and magnitude of the cognate inflationary CD8^+^ T-cell response (18). We used three well-characterized MCMV mutants expressing low-avidity or high-avidity epitopes in different genetic contexts: the recombinant MCMV^ie2SL^, which expresses the high avidity HSV-1 epitope SSIEFARL (SL) inserted at the C terminus of the *immediate-early 2 (ie2)* gene (18); the MCMV^ie2KNL^ mutant expressing the low-avidity epitope KCSRNRQYL (KNL) also inserted at the C terminus of the *ie2* gene (20); and the MCMV^m45SL^ recombinant, which expresses the same epitope as MCMV^ie2SL^ inserted in a different genetic context, the early *M45 gene* (18).

SL-specific and KNL-specific CD8^+^ T-cells were analysed 120 days post infection (dpi) using tetramer staining to determine the magnitude and the phenotype of the inflating T-cell response. As previously described (20), the SL and the KNL epitopes expressed within the *ie2* gene induced larger inflationary T-cell responses than the SL epitope expressed within the *m45* gene (Figure 1A). MCMV^ie2SL^ induced the inflationary response of the highest magnitude; a median of 13% of total CD8^+^ T-cells were SL tetramer-positive. The size of the specific response to MCMV^ie2KNL^ was approximately 9% of the CD8^+^ T-cell pool, and significantly larger than the response to MCMV^m45SL^, which remained below 5%. Yet even the latter recombinant virus induced a clearly detectable tetramer-positive CD8^+^T-cell population (Figure 1A).

**Figure 1.**
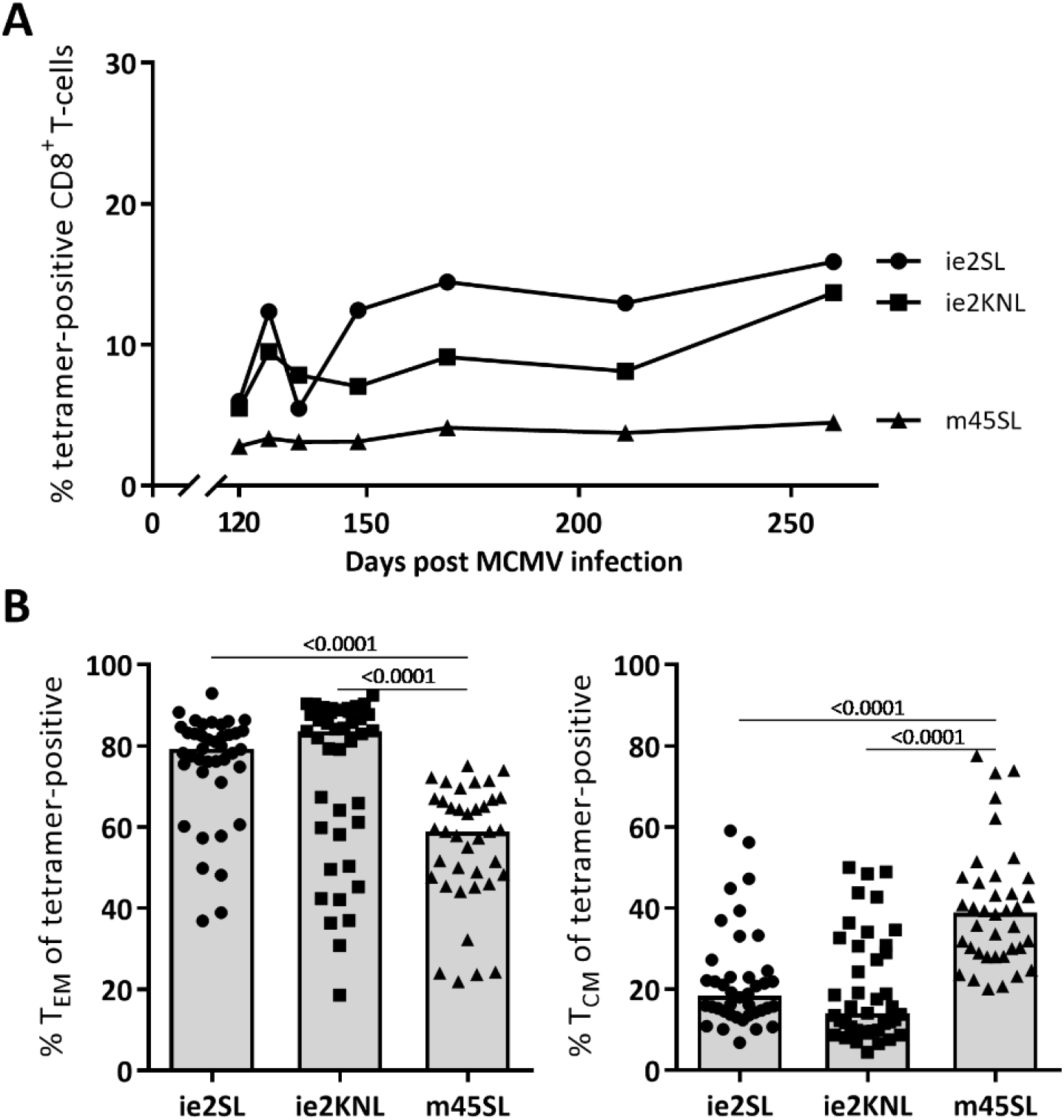
Recombinant viruses induce inflationary responses of different magnitude and phenotype. Mice (129/Sv) were infected with MCMV^ie2SL^, MCMV^ie2KNL^ or MCMV^m45SL^, and 120 days post infection (dpi) tetramer-positive CD8^+^ T-cells from spleen were characterized. **(A)** Median percentage of tetramer-positive cells within CD8^+^ T-cells over time (N=4-7 per time point per group). P-value=0.04 for pooled time points of MCMV^ie2SL^ vs MCMV^ie2KNL^, P-value<0.0001 for pooled time points of MCMV^ie2SL^ vs MCMV^m45SL^, and P-value=0.0006 for pooled time points of MCMV^ie2KNL^ vs MCMV^m45SL^. Pooled samples of different time points were compared using Kruskal-Wallis and Dunn’s multiple comparisons test. Data are pooled from two independent experiments. **(B)**Percentage of T_EM_ (left; CD44^+^CD62L^−^) and T_CM_ (right; CD44^+^CD62L^+^) cells within the tetramer-positive CD8^+^ T-cell pool (MCMV^ie2SL^ N=39, MCMV^ie2KNL^ N=41, MCMV^m45SL^ N=38). Data are pooled from two independent experiments. Bars represent the median percentage. For the % of T_EM_ P-value>0.999 for pooled time points of MCMV^ie2SL^ vs MCMV^ie2KNL^, P-value<0.0001 for pooled time points of MCMV^ie2SL^ vs MCMV^m45SL^, and P-value<0.0001 for pooled time points of MCMV^ie2KNL^ vs MCMV^m45SL^. For the % of T_CM_ P-value=0.976 for pooled time points of MCMV^ie2SL^ vs MCMV^ie2KNL^, P-value<0.0001 for pooled time points of MCMV^ie2SL^ vs MCMV^m45SL^, and P-value<0.0001 for pooled time points of MCMV^ie2KNL^ vs MCMV^m45SL^. Pooled samples of different time points were compared using Kruskal-Wallis and Dunn’s multiple comparisons test. Median percentage of T_N_, T_CM_ and T_EM_ cells within tetramer-negative and tetramer-positive CD8^+^ T-cells in MCMV-infected and uninfected mice are shown in Sup. Figure 3. Changes in the median percentage of T_N_, T_CM_ and T_EM_ CD8^+^ T-cells over time are shown in Sup. Figure 4.

Antigen-specific CD8^+^ T-cells composing an inflationary response typically present an effector phenotype and maintain their effector function (14). Accordingly, the vast majority (>80%) of tetramer-positive CD8^+^ T-cells composing large inflationary responses (MCMV^ie2SL^ and MCMV^ie2KNL^) had a T_EM_ phenotype (CD44^+^CD62L^−^). In contrast, only 60% of the tetramer-positive CD8^+^ T-cells induced by MCMV^m45SL^ presented a T_EM_ phenotype, while the remaining 40% expressed T_CM_ markers (CD44^+^CD62L^+^) (Figure 1B, Sup. Figure 3 and Sup. Figure 4). Less than 2% of the tetramer-positive T-cells had a T_N_ phenotype (CD44^−^CD62L^+^).

### Ki-67 expression pattern of CD8^+^ T-cells does not differ between MCMV-induced inflationary responses of different magnitude

To study the dynamics of CD8^+^ T-cells in the stable phase of chronic MCMV infection, we first determined cell proliferation by measuring Ki-67 expression. The fraction of Ki-67^hi^ cells within T_N_, T_CM_ and T_EM_ CD8^+^ T-cells was not significantly different between uninfected mice and chronically infected mice for all 3 viruses (Figure 2). In line with previous reports (26), we found that the percentage of Ki-67^hi^ cells was the lowest within T_N_ cells (median over all groups of 0.4%), intermediate within T_CM_ cells (median of 4.5%) and the highest within T_EM_ cells (median of 12%) (Figure 2B). Approximately 5% of the tetramer-positive CD8^+^ T-cells were Ki-67^hi^. For each viral infection, the total fraction of Ki-67^hi^ cells within the tetramer-positive cells was not significantly different from that of memory phenotype T-cells, and was in fact between the Ki-67 expression levels of T_EM_ and T_CM_ cells (Figure 2B). Based on Ki-67 expression, we thus found no indication that inflated MCMV-specific CD8^+^ T-cells have different proliferation rates than bulk memory phenotype CD8^+^ T-cells. Since Ki-67 only provides a snapshot marker of T-cell proliferation, we next studied the average production and loss rates of the cells using *in vivo* deuterium labelling.

**Figure 2.**
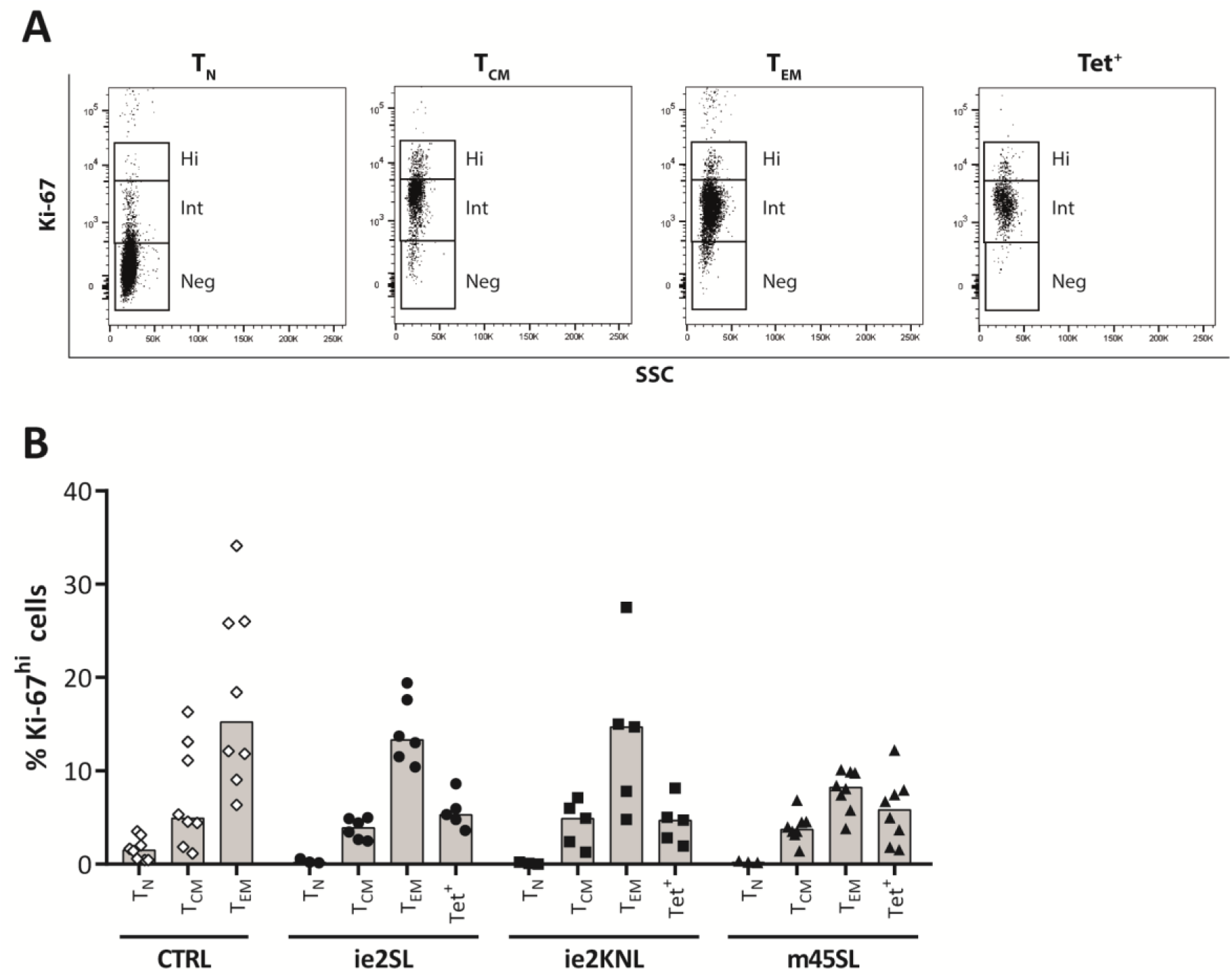
Percentage Ki-67^hi^ cells within CD8^+^ T-cell subsets in MCMV-infected and uninfected mice. **(A)** Representative Ki-67 staining of T_N_, T_CM_ and T_EM_ and Tet^+^ CD8^+^ T-cells from blood of a MCMV^m45SL^-infected mouse 17 months post infection. **(B)**Fraction of Ki-67^hi^ T_N_, T_CM_ and T_EM_ cells and tetramer-positive (Tet^+^) CD8^+^ T-cells from blood of MCMV^ie2SL^ (N=4-6), MCMV^ie2KNL^ (N=4-5) and MCMV^m45SL^ (N=4-8) infected mice and age-matched and sex-matched uninfected mice (CTRL, N=8). For MCMV^ieKNL^ P-value=0.418 for pooled time points of % Ki-67^hi^ Tet^+^ vs T_EM_, P-value>0.999 for pooled time points of % Ki-67^hi^ Tet^+^ vs T_CM_, and P-value=0.224 for pooled time points of % Ki-67^hi^ Tet^+^ vs T_N_; for MCMV^ie2SL^ P-value=0.583 for pooled time points of % Ki-67^hi^ Tet^+^ vs T_EM_, P-value>0.999 for pooled time points of % Ki-67^hi^ Tet^+^ vs T_CM_, and P-value=0.389 for pooled time points of %Ki-67^hi^ Tet^+^ vs T_N_; for MCMV^m45SL^ P-value=0.938 for pooled time points of % Ki-67^hi^ Tet^+^ vs T_EM_, P-value>0.999 for pooled time points of % Ki-67^hi^ Tet^+^ vs T_CM_, and P-value=0.088 for pooled time points of % Ki-67^hi^ Tet^+^vs T_N_. Pooled samples of different time points were compared using Kruskal-Wallis and Dunn’s multiple comparisons test. Data are pooled from two independent experiments. Bars represent median percentages.

### Kinetics of T_N_, T_CM_, and T_EM_ CD8^+^ T-cells during chronic MCMV infection

The *in vivo* kinetics of memory CD8^+^ T-cells have primarily been studied in bulk memory phenotype T-cells (5). Here, we quantified the dynamics of T_N_, T_CM_, T_EM_ and total memory (T_M_) CD8^+^ T-cell subsets in chronically-infected and uninfected 129/Sv mice. Mice received ^2^H_2_O for 4 weeks and were sacrificed at different time points during the labelling and the de-labelling period. We subsequently used previously published mathematical models (5) (see material and methods) to quantify the average production and loss rates of T_N_, T_CM_, T_EM_ and T_M_ CD8^+^ T-cells based on their deuterium labelling data.

Deuterium enrichment curves of T_N_, T_CM_, T_EM_ and T_M_ cells were very similar in MCMV^ie2SL^, MCMV^ie2KNL^, and MCMV^m45SL^ infected animals (Figure 3). In line with this, the best fits of the model to the data yielded similar estimates for the average production rates *p* and the average loss rates of labelled cells *d\** within the T_N_, T_CM_, T_EM_ and T_M_ CD8^+^ T-cell populations for the three different viruses (Figure 4A and Table 1). The estimated average production rate, *p*, of T_N_ cells during chronic MCMV infection was 0.0090 per day for MCMV^ie2SL^, 0.0095 per day for MCMV^ie2KNL^, and 0.0077 per day for MCMV^m45SL^, suggesting that T_N_ cells turn over relatively little. In contrast, T_CM_ cells turned over significantly, with average production rates of 0.0132 per day for MCMV^ie2SL^, 0.0151 per day for MCMV^ie2KNL^, and 0.0139 per day for MCMV^m45SL^. The average production rates of T_EM_ cells were consistently the highest, with 0.0204 per day for MCMV^ie2SL^, 0.0229 per day for MCMV^ie2KNL^, and 0.0205 per day for MCMV^m45SL^ (Table 1).

**Figure 3.**
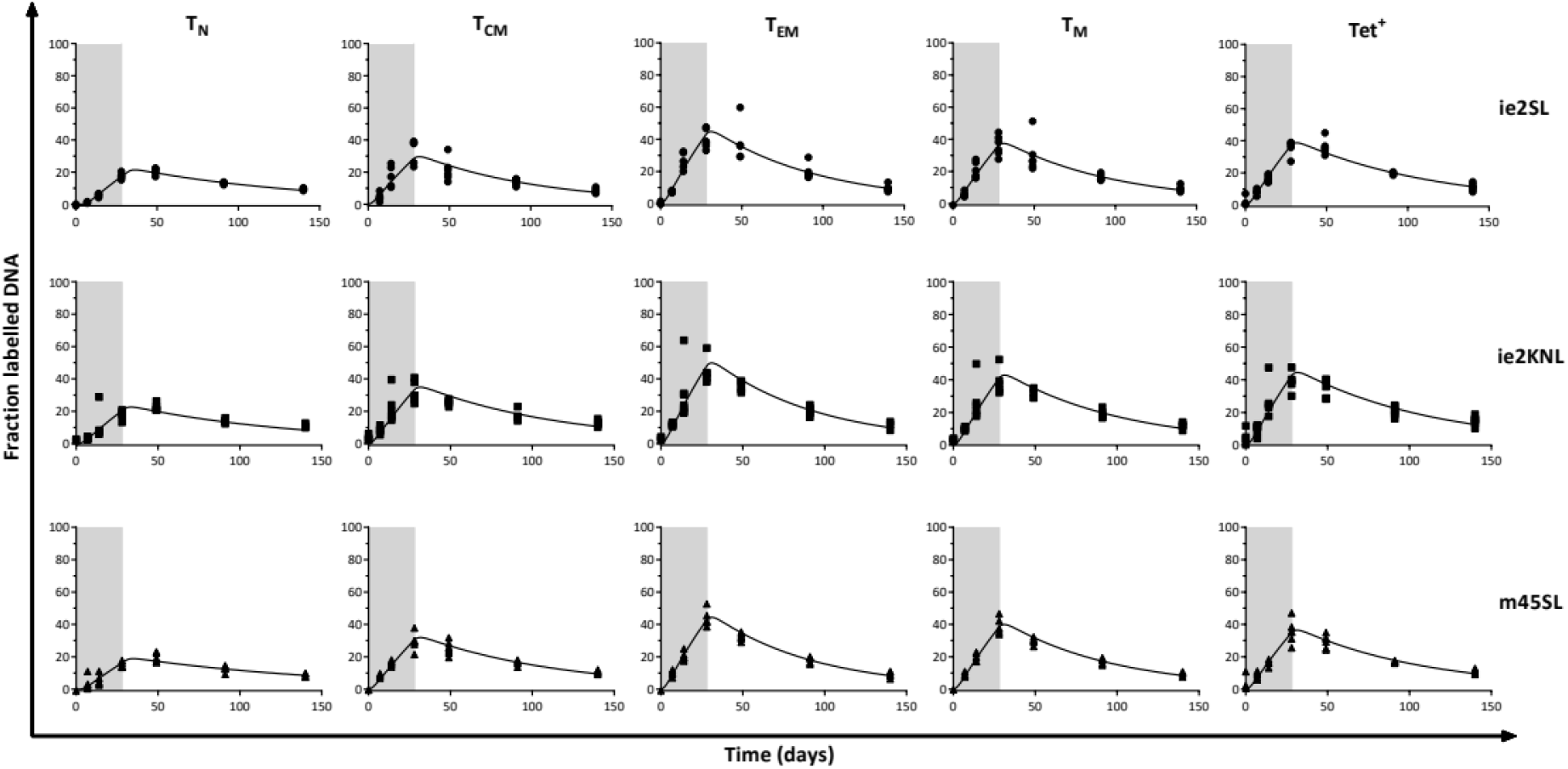
Deuterium labelling of tetramer-negative and tetramer-positive CD8^+^ T-cells in MCMV-infected mice. Deuterium enrichment in the DNA of T_N_, T_CM_, T_EM_ and T_M_ and tetramer-positive (Tet^+^) CD8^+^ T-cells 120 days after MCMV^ie2SL^, MCMV^ie2KNL^ or MCMV^m45SL^ infection. The curves represent the best fits of the model (5) to the deuterium enrichment data. Label enrichment was scaled between 0 and 100% by dividing all enrichment levels by the estimated maximum enrichment level of thymocytes (Sup. Figure 2 and Sup. Table 1). Parameter estimates corresponding to the best fits are given in Table 1.

**Figure 4.**
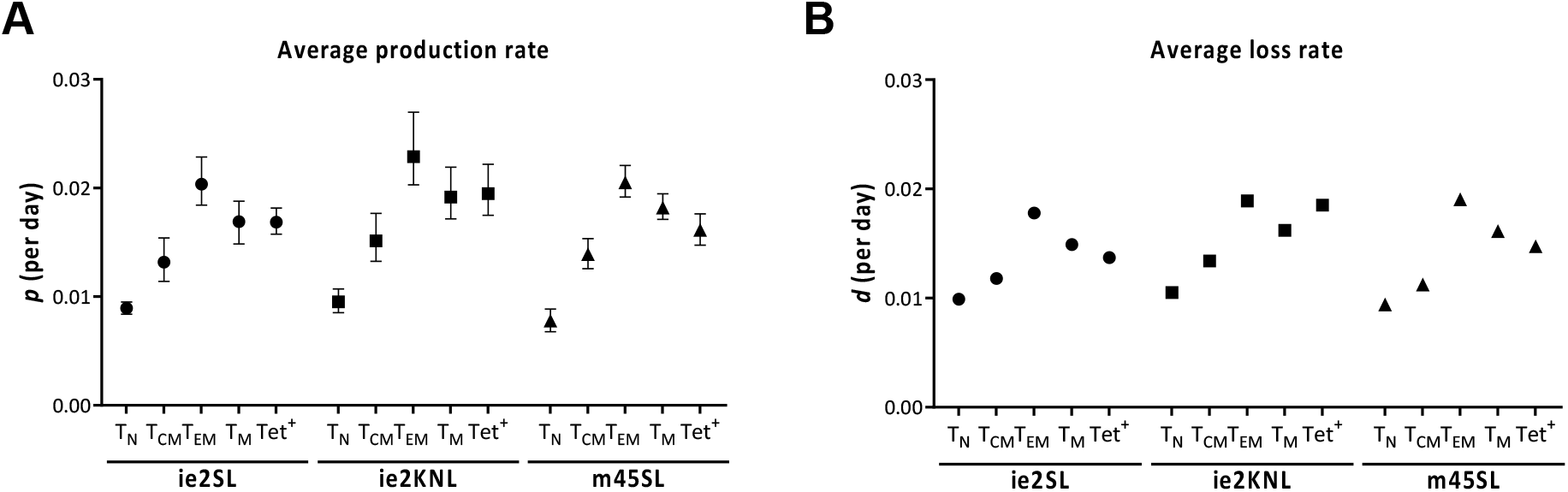
Average production and loss rates of T-cells in MCMV-infected mice. Estimated average production rates *p* per day (A) and average loss rates *d* per day (B) for T_N_, T_CM_, T_EM_ and T_M_ CD8^+^ T-cells and tetramer-positive (Tet^+^) memory CD8^+^ T-cells. (A) Average production rates *p* were based on the best fits of the deuterium labelling data of Figure 3. Their values are reported in Table 1. Whiskers represent the corresponding 95% confidence intervals. (B) The average loss rates *d* were calculated from *d* = *p* – *r*, using the best estimates of *p* (Table 1) and the estimated growth rate *r* of the specific T-cell population (Table 2). Because the average loss rates *d* were calculated based on other parameters, they are given without 95% confidence intervals.

### The turnover rate of MCMV-specific CD8^+^ T-cells is independent of the magnitude of the inflationary response

To investigate how the size of an MCMV-specific memory T-cell response is related to its turnover (19), we quantified the turnover rates of Tet^+^ CD8^+^ T-cells composing large (MCMV^ie2SL^) and intermediate (MCMV^ie2KNL^) inflationary responses, as well as a low-inflationary response (MCMV^m45SL^). Despite the clear differences in the height of the antigen-specific responses induced by these three viruses, the corresponding deuterium enrichment curves of Tet^+^ CD8^+^ T-cells were very similar (Figure 3). The best fits of the model to the deuterium-enrichment data of SL-tetramer-positive CD8^+^ T-cells in MCMV^ie2SL^ and MCMV^m45SL^ infected mice (Figure 3) confirmed that, despite the 3-fold difference in the size of these inflationary responses (Figure 1A) and their different T_CM_ and T_EM_ compositions (Figure 1B), their average production rates *p* and loss rates of labelled cells *d\** were not significantly different (Figure 4A and Table 1). We estimated that SL tetramer-positive CD8^+^ T-cells had an average production rate of 0.0169 per day in MCMV^ie2SL^ infected mice and 0.0161 per day in MCMV^m45SL^ infected mice. For KNL-specific T-cells, which comprised the intermediate inflationary response, we found an average production rate of 0.0195 per day (Table 1). Thus, there were no substantial differences in the average turnover rates of tetramer-positive CD8^+^ T-cells composing large, intermediate and low inflationary responses.

### MCMV-specific CD8^+^ T-cells do not have significantly longer lifespans than memory phenotype CD8^+^ T-cells

Finally, in order to investigate the hypothesis that accumulation of inflationary responses in MCMV is due to accumulation of long-lived cells, we compared the average turnover rates of MCMV-specific (Tet^+^) CD8^+^ T-cells to those of tetramer-negative total memory (T_M_) CD8^+^ T-cells (see Methods). When comparing the best fits of the individual datasets, we found no statistical indication that TM and Tet^+^ T cells had different production rates *p* or loss rates of labelled cells *d\** (Figure 4A and Table 1).

Since the average loss rate *d\** of labelled cells may not be representative of the cell population as a whole (23), we used additional information on absolute cell numbers to compare the average loss rates *d* of Tet^+^ and T_M_ cells. Although these absolute cell numbers are notoriously noisy, we estimated a slight increase in cell numbers in the T_CM_, T_EM_, T_M_ and tetramer-positive CD8^+^ T-cell populations (Sup. Figure 5 and Table 2). The average production rates *p* may thus not be equal to the average loss rates *d* of cells. Even when accounting for this rate of increase, *r*, in cell numbers, we found very similar average loss rates *d* (where *d* = *p* – *r*, see Methods) of T_M_ and Tet^+^ cells in MCMV^ie2SL^, MCMV^ie2KNL^ and MCMV^m45SL^ infected mice (Figure 4B and Table 2). We thus found no evidence for the previously proposed idea that CMV-specific T-cells are longer-lived than other memory T-cells (27).

Although an advantage of the present study is the face-to-face comparison of tetramer-positive and tetramer-negative cells in the same mouse, it is more than likely that the tetramer-negative T-cell populations still contained MCMV-specific CD8^+^ T-cells specific for other MCMV-epitopes (17, 18, 20). We wondered whether this could have masked possible differences in the turnover of MCMV-specific and non-MCMV-specific memory CD8^+^ T-cells. To investigate this, we compared the deuterium enrichment levels of T_N_, T_CM_, T_EM_ and T_M_ cells in MCMV-infected mice to those in uninfected mice. Since these levels were very similar (Sup. Figure 6), we conclude that the expected lifespan of MCMV-specific memory CD8^+^ T-cells is not significantly different from that of other memory-phenotype CD8^+^ T-cells.

## Discussion

During the chronic phase of MCMV infection, we found no evidence that MCMV-specific CD8^+^T-cells are longer-lived or produced at higher rates than bulk memory-phenotype CD8^+^ T-cells. These findings are in line with our recent findings in humans, which showed that CMV-specific CD8^+^ T-cells had similar turnover rates as bulk memory CD8^+^ T-cells (Van den Berg et al. submitted). Both outcomes are remarkable in the light of a previously published deuterated-glucose labelling study in humans, which reported that CMV-specific CD8^+^ T-cells incorporated less deuterium than CD45RO^+^ (memory) T-cells (19), which led to the hypothesis that inflating responses are composed of relatively long-lived memory T cells.

We found that the turnover rates of antigen-specific T-cells composing inflationary responses that varied up to 3-fold in size were not significantly different. This adds further support to our conclusion that the magnitude of inflationary responses is not explained by extended lifespans of MCMV-specific T-cells. Instead, our data suggest that the explanation for the size differences between MCMV-specific CD8^+^ T-cell responses and for memory inflation in general should be sought earlier during infection. In line with this, it has recently been shown that the inflationary potential of CMV-specific T-cells is set early, during the acute phase of the response, and is reflected by their early transcriptomic profile (28).

To interpret the deuterium labelling data, we used a previously proposed kinetic heterogeneity model (23), which yields the average production rate *p* of cells, as well the average loss rate *d\** of labelled cells. It was previously explained that cell populations in steady state typically yield *d\** > *p*, because *p* is representative of all cells in the population, while *d\** is biased towards cells that have just divided (23). The estimated value of *p* can thus safely be interpreted as the average production rate of cells. Assuming that the vast majority of MCMV-specific cells are formed by peripheral T-cell division, and not by continuous recruitment of new naive MCMV-specific T-cells into the memory pool, the similar production rates of Tet^+^ and T_M_ cells imply that MCMV-specific T-cells do not divide more frequently than other memory T-cells, which is supported by their similar Ki67-expression levels. To compare the average loss rates of Tet^+^ and T_M_ cells, we used additional information on absolute cell numbers, which confirmed that also the expected lifespans of Tet^+^ and T_M_ cells are very similar. We thus found no evidence for the previously proposed idea that CMV-specific T-cells are longer-lived than other memory T-cells (27).

Non-steady cell numbers in some of the populations may explain why we sometimes found values of *d\** lower than *p* (see Table 1). This is typically not observed in deuterium labelling experiments (23), and suggests that cells that have recently divided live longer than other cells in the population. Alternatively, for populations that are not in steady state, if cellular turnover is dependent on cell densities, average production rates may decrease during the labelling experiment, while cell numbers are increasing. This could explain why, contrary to what is typically observed for populations in steady state, some turnover rates during the de-labelling phase were lower than during the up-labelling phase (Swain et al. manuscript in preparation).

A previous study in humans reported that YFV-specific CD8^+^ T-cells triggered upon YFV vaccination divide sporadically, approximately every 666 days (11), and less than bulk memory T-cells. It was argued that the shorter intermitotic times of bulk memory T-cells probably reflect their continuous antigen stimulation (11). In line with this hypothesis, we found that MCMV-specific CD8^+^T-cells, which are repeatedly exposed to their cognate antigen, have similar lifespans as bulk memory-phenotype CD8^+^ T cells, which may also be continuously exposed to commensal and environmental antigens. However, the observation that lymphocytic choriomeningitis virus (LCMV)-specific memory CD8^+^T-cells transferred into naive mice had similar turnover rates as bulk memory-phenotype CD8^+^ T-cells (29), suggests that even antigen-specific T-cells maintained in the absence of cognate antigen can turnover as fast as bulk memory-phenotype T-cells. Although we cannot exclude the possibility that the differences in the maintenance of YFV-, MCMV- and LCMV-specific memory CD8^+^ T-cells are due to mouse-man differences, the characteristics of different antigen-specific memory T-cells may also depend on the nature of the infection, the duration of the stimulus, and the concomitant response to other antigens. It is therefore perhaps not surprising that antigen-specific T-cell responses against different infections have different dynamics (30). Future studies into the dynamics of memory T-cells specific for antigens that are presented persistently (chronic), intermittently (latent reactivating) or only once (acute) are needed to gain more insight into how antigen-specific memory T-cell responses are maintained in mice and man.

Chronic CMV infection in both mice and humans is under constant immune surveillance and triggers ongoing CD8^+^ T-cell responses. It is thought that CMV infection modulates the peripheral lymphoid pool (31, 32), and affects T-cell differentiation and function (33), not only of CMV-specific T-cells but also of T-cells with other specificities (34). Under this hypothesis, we directly compared the dynamics of T_N_, T_CM_, T_EM_ and T_M_ cells in uninfected and chronically MCMV-infected mice [Sup. Figure 6], and found no significant differences in their kinetics. Despite differences in the composition of the T-cell pool, our results therefore suggest that the dynamics of non-MCMV specific CD8^+^ T-cells are not substantially affected during chronic MCMV infection. We previously observed that also cellular immune function was maintained during latency, as responses to heterologous virus infection and immune protection were not diminished in mice latently infected with MCMV or other herpesviruses (35).

The large prevalence of chronic CMV infection in the human population (>50%) (36) and its effect on healthy aging (37–39), together with the emerging interest in CMV-based vector vaccines (15), highlight the need to understand how CMV-specific CD8^+^ T-cell responses are maintained. *In vivo* MCMV infection provided us with the means to address fundamental questions about the maintenance and turnover of inflated CD8^+^ T-cell responses. The finding that the maintenance of inflationary MCMV-specific CD8^+^ T-cells does not differ from that of low-inflationary memory CD8^+^ T-cells suggests that inflationary CD8^+^ T-cell responses, such as those induced by CMV-based vector vaccines, may also result in a memory CD8^+^ T-cell response of high magnitude without substantial alterations in the dynamics of the cells.

## Acknowledgments

We thank Ramon Arens for generously providing all MHC-peptide tetramers, Nienke Vrisekoop for help and support with the animal experiments, Derek Macallan and Linda Hadcocks for measuring ^2^H_2_O enrichment in the body water of the uninfected mice, and Janine Schreiber and Lothar Gröbe for technical assistance.

