## Supplementary material for "Turnover of MCMV-expanded CD8^+^ T-cells is similar to that of memory phenotype T-cells and independent of the magnitude of the response"

### Supplementary figures

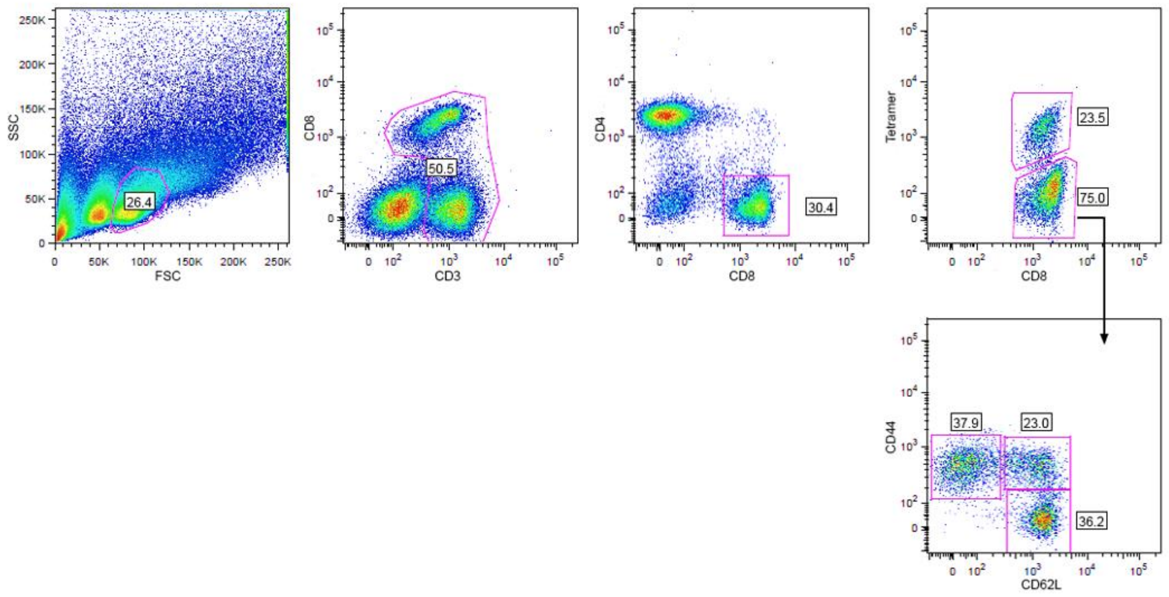


**Sup. Figure 1. Sort gating strategy.** Strategy for sorting of tetramer-positive $CD8^+$ T cells and tetramer-negative $T_N$ ($CD62L^+CD44^-$), $T_{CM}$ ($CD62L^+CD44^+$) and $T_{EM}$ ($CD62L^-CD44^+$) $CD8^+$ T-cells.

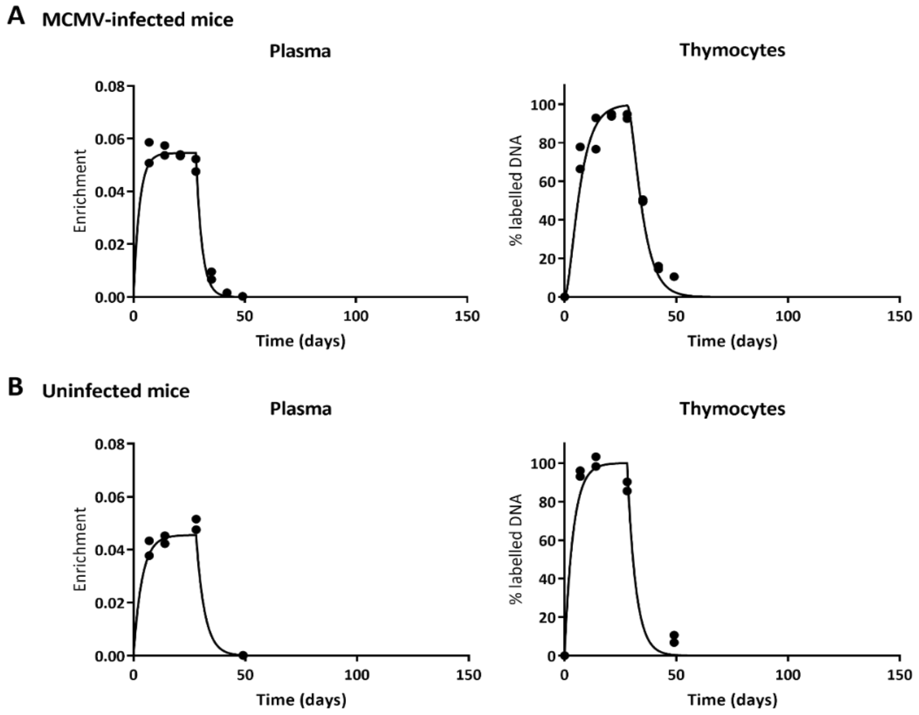


**Sup. Figure 2. Deuterium enrichment in plasma and thymocytes.** Deuterium enrichment in the body water (plasma) and thymocytes. Symbols represent individual measurements during the up- and down-labelling phases of **(A)** MCMV-infected mice and **(B)** age-matched and sex-matched uninfected mice. The labeling data of thymocytes were fitted with Equation (2) assuming $p = d^*$, and their label enrichment was scaled between 0 and 100% by normalizing for their estimated maximum level of deuterium enrichment. The enrichment curves of thymocytes were used to normalize the deuterium enrichment data of the different T-cell subsets (see material and methods). Parameter estimates of the best fits are given in Sup. Table 1.

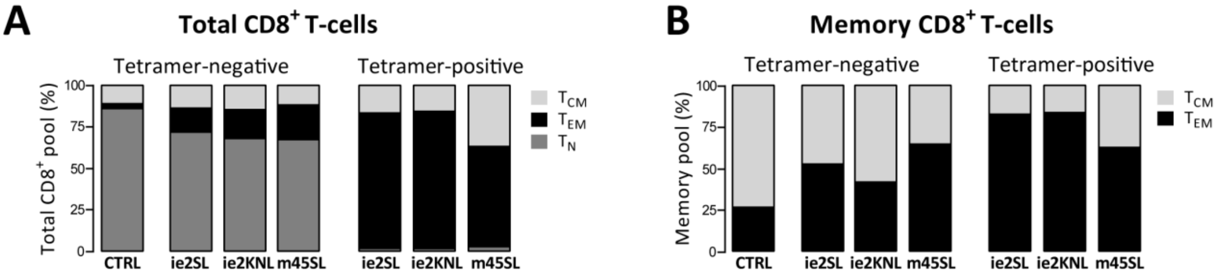


**Sup. Figure 3. Composition of the CD8$^+$ T-cell pool in MCMV-infected and uninfected mice. (A)** Median percentage of $T_N$, $T_{CM}$ and $T_{EM}$ cells within tetramer-negative and tetramer-positive total CD8$^+$ T-cells and **(B)** median percentage of $T_{CM}$ and $T_{EM}$ cells within tetramer-negative and tetramer-positive memory phenotype CD8$^+$ T-cells (CD44$^+$) from MCMV-infected mice (MCMV$^{ie2SL}$ N=39, MCMV$^{ie2KNL}$ N=41, MCMV$^{m45SL}$ N=38) and age- and sex-matched uninfected mice (CTRL N=10).

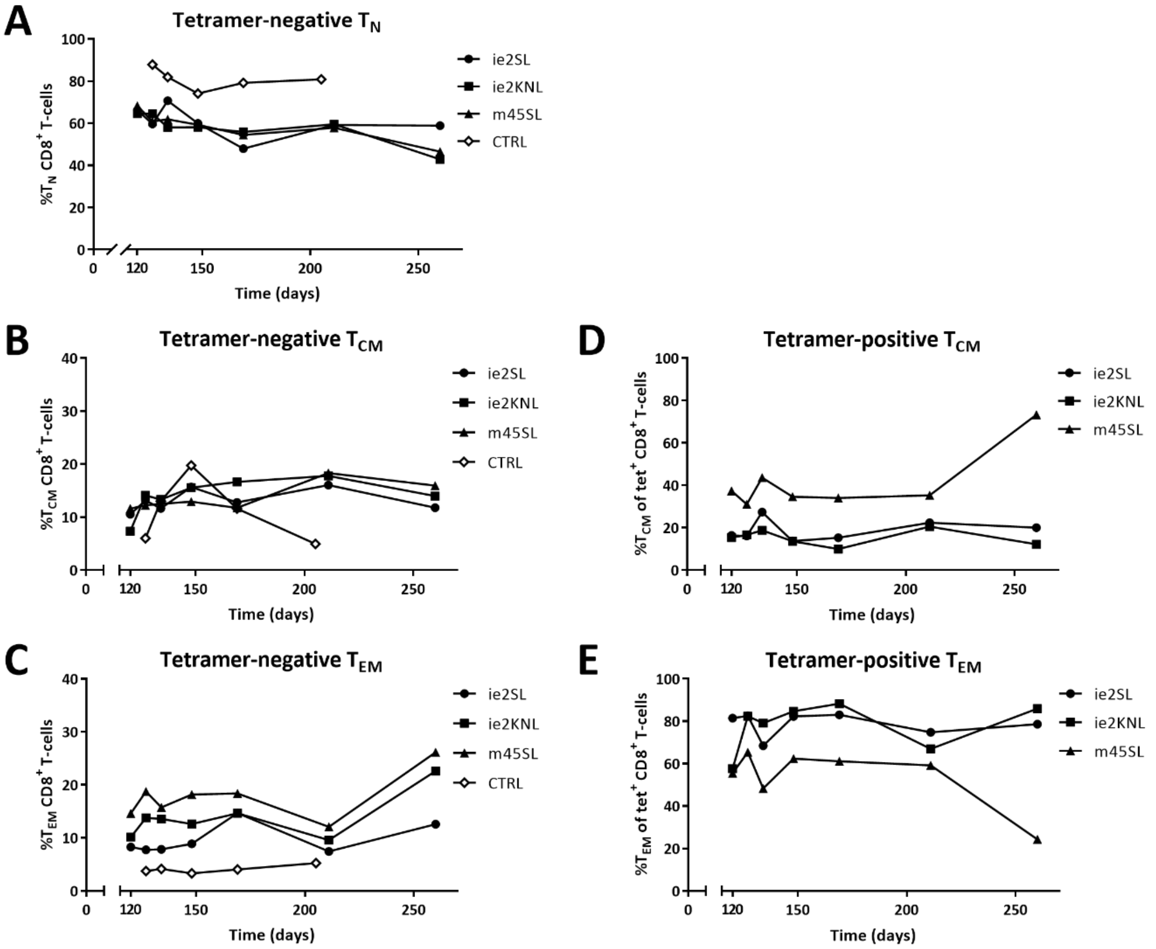


**Sup. Figure 4. Composition of the $CD8^+$ T-cell pool over time in MCMV-infected and uninfected mice.** Mice (129/Sv) were infected at t=0 with $MCMV^{ie2SL}$, $MCMV^{ie2KNL}$ or $MCMV^{m45SL}$, age- and sex-matched uninfected 129/Sv mice were used as controls (CTRL). From 120 dpi onwards, tetramer-negative and tetramer-positive $CD8^+$ T-cells from spleen were characterized. Median percentage of **(A)** $T_N$, **(B)** $T_{CM}$ and **(C)** $T_{EM}$ cells within tetramer-negative $CD8^+$ T-cells and median percentage of **(D)** $T_{CM}$ and **(E)** $T_{EM}$ cells within tetramer-positive $CD8^+$ T-cells over time (for MCMV-infected mice, N=4-7 per time point; for CTRL mice, N=2 per time point).

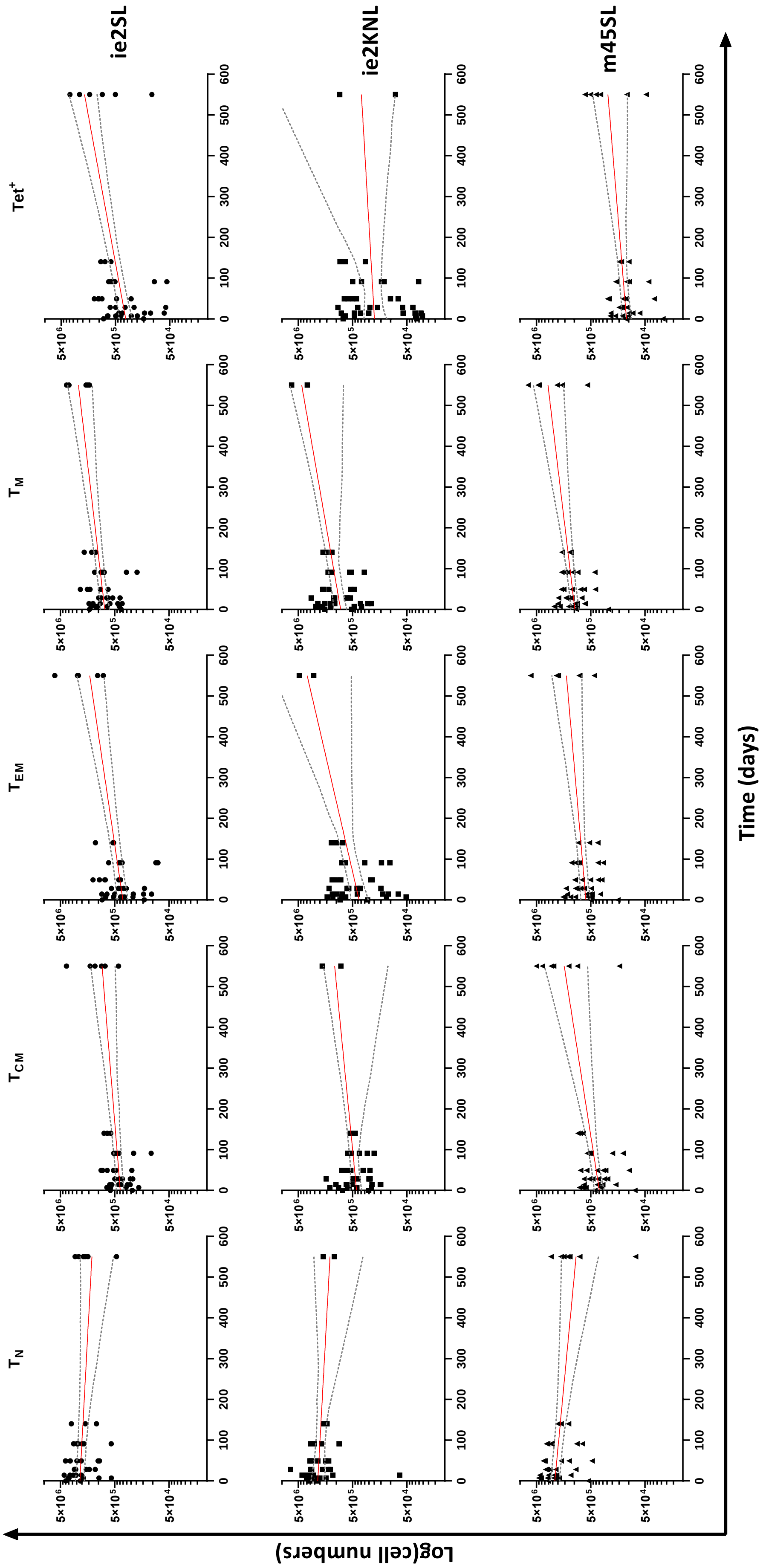

T_N
T_CM
T_EM
T_M
Tet+
ie2SL
ie2KNL
m45SL
5×10^6
5×10^5
5×10^4
0
100
200
300
400
500
600
Log(cell numbers)
Time (days)

**Sup. Figure 5. Estimation of the growth/decay rate *r* of the different T-cell populations.** We estimated the rate of change in cell numbers, $r$, and the cell numbers at the start of the experiment (i.e. 120 days p.i.), *N*(0), in the $T_N$, $T_{CM}$, $T_{EM}$, $T_M$, and $Tet^+$ T-cell populations of MCMV-infected mice by fitting a simple exponential growth/decay model to the observed cell numbers in mice over 550 days (see Methods). The best fits to the data are given by the red curves. The grey curves represent the 95% confidence intervals (CI) and were calculated by taking the 95% CI, at each time point, of 500 bootstrap trajectories (see Methods). The estimated values of *r* and *N*(0) and their 95% CI are given in Table 2 and Sup. Table 2, respectively.

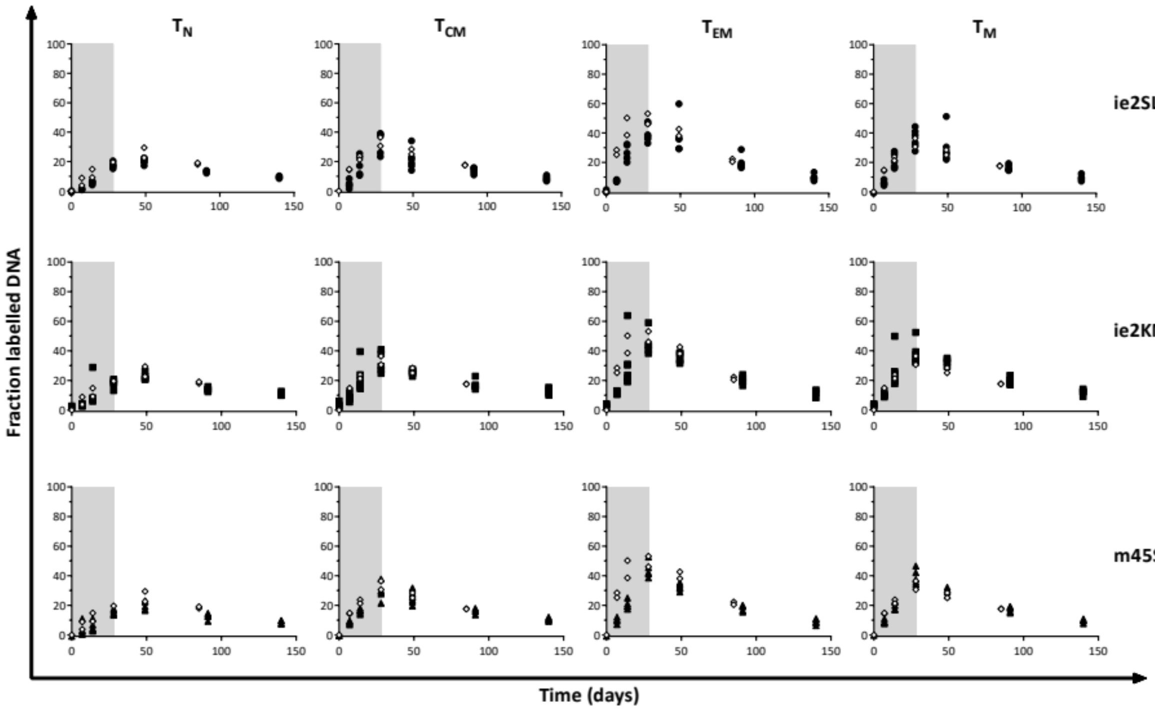


**Sup. Figure 6. Enrichment level in $T_N$, $T_{CM}$, $T_{EM}$ and $T_M$ $CD8^+$ T-cells of uninfected mice is similar to that of MCMV-infected mice.** $^2H$ enrichment in the DNA of $T_N$, $T_{CM}$, $T_{EM}$ and $T_M$ of uninfected 129/Sv mice (open diamonds) and tetramer-negative $T_N$, $T_{CM}$, $T_{EM}$ and $T_M$ $CD8^+$ T-cells of $MCMV^{ie2SL}$, $MCMV^{ie2KNL}$ or $MCMV^{m45SL}$ infected mice (black symbols, which are also shown in Figure 3).

**Sup. Table 1. Estimated parameters and their corresponding 95% confidence intervals for deuterium enrichment in plasma and thymocytes of MCMV-infected and uninfected mice.** Since allowing for different values of $p$ and $d^*$ did not significantly improve the description of the labeling curve of the thymocytes, we report the best fits of the model with $p = d^*$. For uninfected mice, the turnover rate of thymocytes was non-identifiable (N.I.).

| | | MCMV-infected mice | Uninfected mice |
|---|---|---|---|
| Plasma | $\delta$ (per day) | 0.3730 (0.2436;0.3844) | 0.2616 (0.1945;0.4198) |
| | $f$ (per day) | 0.0545 (0.0526;0.0829) | 0.0455 (0.0430;0.0483) |
| | $S_0$ | 0.0006 (0;0.0007) | 0 (0;0) |
| Thymocytes | $c$ | 3.0271 (2.7906;3.2116) | 3.9011 (3.5946;4.1589) |
| | $p$ | 0.1839 (0.1507;0.2314) | N.I. |

**Sup. Table 2. Estimated values of *N*(0).** Overall growth/decay rates *r* of the specific T-cell populations (Sup. Figure 5 and Table 2) were estimated by simultaneously estimating each population size at the start of the experiment (i.e. 120 days p.i.), *N*(0), of which the values are given below, with 95% CI in brackets.

| | Estimated population size at start of experiment, *N*(0) | | |
|---|---|---|---|
| $CD8^+$ T-cell subset | $MCMV^{ie2SL}$ | $MCMV^{ie2KNL}$ | $MCMV^{m45SL}$ |
| $T_N$ | 2187085 (1800056; 2643753) | 2193015 (1597846; 2908849) | 2286946 (1855345; 2810501) |
| $T_{CM}$ | 394773 (333631; 467489) | 423450 (341455; 559050) | 346660 (283320; 431499) |
| $T_{EM}$ | 353468 (286233; 451371) | 380288 (258420; 541734) | 614460 (509244; 760951) |
| $T_M$ | 763465 (624839; 918752) | 832600 (643671; 1113140) | 979134 (819990; 1168923) |
| $Tet^+$ | 324228 (246843; 424190) | 198686 (117330; 308198) | 109771 (89367; 132410) |
